# H_2_S preconditioning induces long-lived perturbations in O_2_ metabolism

**DOI:** 10.1101/2023.10.20.563353

**Authors:** David A. Hanna, Jutta Diessl, Arkajit Guha, Roshan Kumar, Anthony Andren, Costas Lyssiotis, Ruma Banerjee

**Affiliations:** Departments of Biological Chemistry, University of Michigan Medical Center, Ann Arbor, MI 48109-0600; Departments of Molecular and Integrative Physiology, University of Michigan Medical Center, Ann Arbor, MI 48109-0600; Departments of Internal Medicine, University of Michigan Medical Center, Ann Arbor, MI 48109-0600; Departments of Rogel Cancer Center, University of Michigan Medical Center, Ann Arbor, MI 48109-0600

## Abstract

Hydrogen sulfide exposure in moderate doses can induce profound but reversible hypometabolism in mammals. At a cellular level, H_2_S inhibits the electron transport chain (ETC), augments aerobic glycolysis, and glutamine-dependent carbon utilization via reductive carboxylation; however, the durability of these changes is unknown. We report that despite its volatility, H_2_S preconditioning increases *P*_50(O2)_, the O_2_ pressure for half maximal cellular respiration, and has pleiotropic effects on oxidative metabolism that persist up to 24-48 h later. Notably, cyanide, another complex IV inhibitor, does not induce this type of metabolic memory. Sulfide-mediated prolonged fractional inhibition of complex IV by H_2_S is modulated by sulfide quinone oxidoreductase, which commits sulfide to oxidative catabolism. Since induced hypometabolism can be beneficial in disease settings that involve insufficient or interrupted blood flow, our study has important implications for attenuating reperfusion-induced ischemic injury, and/or prolonging shelf life of biologics like platelets.

---

Clinical outcomes of traumatic injury or diseases that result from insufficient or interrupted blood supply can be improved by decreasing metabolic demand^1, 2^. The discovery of H_2_S as a signaling molecule^3^ paved a paradigm shift in our understanding of its biology from that of a mere environmental toxin to a modulator of mammalian energy metabolism^4, 5^. Moderate exposure to H_2_S (∼80 ppm, 6 h) in mice elicits profound hypometabolism that is characterized by a drop in metabolic rate by 90% and in the core body temperature to 15 °C, which is rapidly reversed in room air^6^. While the mechanism of sulfide-induced hypometabolism remains elusive, an ever-increasing span of physiological effects is ascribed to H_2_S, which encompass major organ systems, including the cardiovascular, gastrointestinal, and central nervous system^7–11^. At a cellular level, the durability and multifaceted implications of “buying time” by suppressing oxidative metabolism, are poorly understood, and the therapeutic promise of sulfide in clinical and experimental models of injury and disease, remains to be realized.

The steady-state levels of H_2_S are estimated to be in the tens of nanomolar range in most tissues^12–14^. Fluctuations in sulfur metabolism due to factors such as dietary sulfur intake, antibiotic use, gut microbial composition, or hypoxia, can potentially disturb the delicate balance between H_2_S synthesis and clearance, allowing cellular levels to spike transiently. In the gut, sulfide-producing and using microbes contribute to colonic concentrations estimated to range from 0.2-2.4 mM H_2_S^15, 16^. Sulfide is efficiently oxidized by sulfide quinone oxidoreductase (SQOR), an inner mitochondrial membrane enzyme^17, 18^, which shields complex IV from respiratory poisoning^19^. The reversible inhibition of complex IV by H_2_S is accompanied by induction of reductive stress, which propagates from the mitochondrion to other compartments, and serves as an important mechanism of H_2_S signaling^4, 5^.

The sulfide oxidation pathway converts H_2_S through a series of oxidation and sulfur transferase steps to thiosulfate and sulfate (Fig. 1A). SQOR commits H_2_S to catabolism, utilizes coenzyme Q (CoQ) as an electron acceptor, and thus resides at the intersection of sulfide oxidation and the electron transport chain (ETC)^20, 21^. SQOR-dependent H_2_S oxidation not only clears this respiratory poison, but also enhances proton-coupled electron transfer, fueling ATP synthesis^19, 22^. However, when concentrations exceed SQOR clearance capacity, H_2_S inhibits complex IV^23^, which manifests as a decrease in the cellular oxygen consumption rate (OCR). The twin effects of H_2_S as a respiratory substrate and an inhibitor influences the redox state of CoQ, which is used by multiple feeders into the ETC. As H_2_S levels rise and complex IV is inhibited, the restricted availability of CoQ causes a reductive shift in the NAD(P)^+^ pool^19^ (Fig. 1A). The ripple effects of the reductive shift in the ETC influence both central carbon and lipid metabolism^19, 24–26^. The interaction between H_2_S and the ETC is also postulated to underlie the profound but reversible ability of H_2_S to trigger hypometabolism, leading to a state of suspended animation even in a non-hibernating animal^6^. Additionally, short-term H_2_S exposure protects mice against lethal hypoxia, suggesting that an H_2_S-dependent decrease in O_2_ metabolism is important for attenuating hypoxia-induced damage^27^.

**Figure 1.**
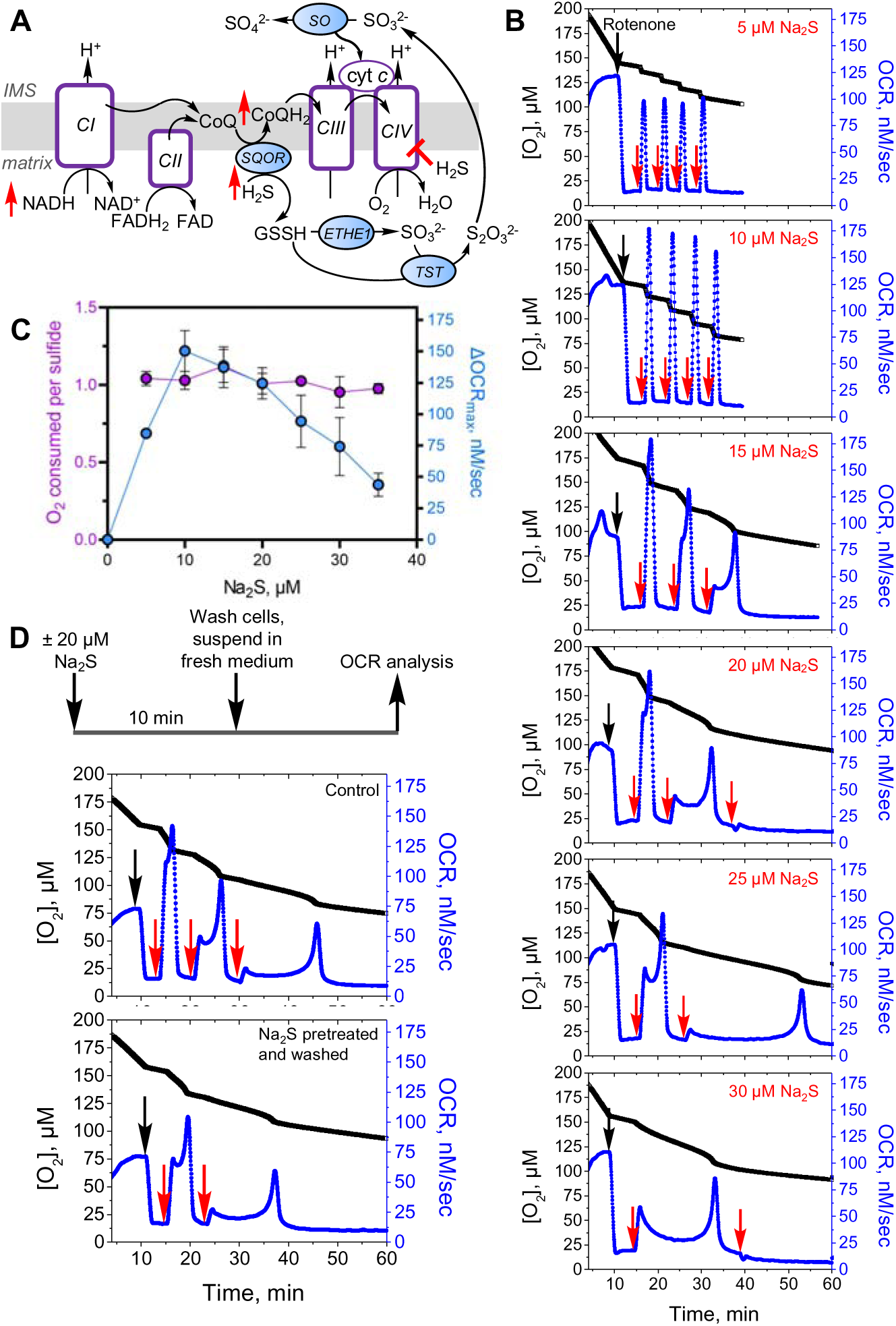
Effect of repeated sulfide exposure on ETC flux. **A**. Sulfide inhibition of complex IV (CIV) causes electron acceptor insufficiency in the ETC due to a predicted build-up of NADH, CoQH_2_, and reduced cytochrome C (cyt C). IMS is intermembrane space, ETHE1 is persulfide dioxygenase, TST is thiosulfate sulfur transferase and SO is sulfite oxidase. **B**. Rotenone (0.5 µM)-treated HT29 cells exposed to low sulfide (5-10 µM) showed sharp increases in OCR with each aliquot of Na_2_S. At higher Na_2_S concentrations (15-30 µM), complex kinetics were observed. **C**. Replotting the data in B as described under Methods, showed a bell-shaped dependence of OCR on sulfide dose and an O_2_:sulfide ratio of 1.0 ± 0.1 between 5-35 µM Na_2_S. The data are representative of at least 4 independent experiments and represent the mean ± SD. **D**. Experimental setup for removing residual sulfide and secreted oxidation products (top). Washing did not impact the persistence of Na_2_S-triggered OCR (bottom). Data are representative of at least 3 independent experiments. Black and red arrows indicate when rotenone and sulfide, respectively were added.

In this study, we report that H_2_S preconditioning elicits persistent bioenergetic perturbations, which we refer to as “H_2_S memory”. The sustained metabolic alterations include an increase in the O_2_ pressure for half maximal respiration (*P*_50(O2)_), and enhanced sensitivity to subsequent H_2_S exposure. The duration of this cellular memory exhibits a dose dependence on H_2_S, is attenuated by dissipation of the mitochondrial NADH pool, but exacerbated by attenuation of SQOR. Furthermore, SQOR expression levels across cell lines correlate with the persistence of H_2_S memory. Collectively, these data reveal that the mechanism underlying the sustained and pleiotropic effects of H_2_S on metabolism, including a reductive shift in cofactor pools, increased aerobic glycolysis, utilization of glutamine via reductive carboxylation and lipid accumulation, as well as changes in the pentose phosphate and pyrimidine biosynthetic pathways. These alterations result from the H_2_S-dependent fractional inhibition of complex IV, revealing a heretofore unappreciated cellular strategy for long-lived regulation of ETC flux.

## RESULTS

### Sulfide elicits dose-dependent changes in oxygen consumption kinetics

Low sulfide concentrations (5-10 µM) elicited sharp increases in O_2_ consumption in HT29 cells, while mixed kinetics were seen at 20 µM Na_2_S (Fig S1A). The different phases at 20 µM Na_2_S presumably represent the sum of OCR activation and inhibition due to the dual action of H_2_S at this concentration. At ≥30 µM Na_2_S, a net decrease in OCR was observed, consistent with respiratory inhibition (Fig. S1B). The protracted recovery times for establishing a new stationary OCR and the persistent fractional inhibition of OCR during the experimental time frame, correlated with sulfide concentration (Fig. S1C,D).

### Effect of repeated acute sulfide exposure on ETC flux

The impact of repeated sulfide addition on OCR was more readily observed in the presence of rotenone, a complex I inhibitor that decreases competition for the CoQ pool (Fig 1B). Under these conditions, the first injection of sulfide generally induced a net increase in OCR, although the kinetics of the response varied in a dose-dependent way. Low sulfide concentrations (<15 µM) triggered a sharp increase followed by a return to basal OCR while higher concentrations (≥20 µM) elicited complex kinetics. At low concentrations (≤10 µM), consecutive sulfide injections led to its clearance with similar kinetics, while at higher concentrations (≥15 µM), complex kinetics and longer clearance times were observed, consistent with sustained inhibition of complex IV.

The maximal change in OCR exhibited a bell-shaped dependence on sulfide concentration, and a stoichiometry of 1.0 ± 0.1 O_2_ consumed to sulfide added was observed across the Na_2_S concentration range (Fig. 1C). Washing cells to remove thiosulfate, a product of the sulfide oxidation pathway that accumulates in the conditioned medium^22^, did not affect sensitivity to subsequent sulfide injections (Fig 1D).

### SQOR increases IC_50_ for complex IV inhibition by H_2_S

Sulfide inhibits complex IV *in vitro* with an estimated *K*_i_ of 0.2 µM with *k*_on_ and *k*_off_ values of 1.5 × 10^4^ M^−1^s^−1^ and 6 x 10^-4^ s^-1^, respectively at 30°C^28^. Sulfide can bind to Cu_B_ (in the 1+ or 2+ oxidation state) and the heme a_3_ site in the MT-CO1 subunit of complex IV (Fig. 2A). We evaluated the magnitude of protection conferred by SQOR by comparing the sulfide-dependent inhibition of OCR in SQOR knockdown (HT29^SQOR^ ^KD^) versus scrambled control (HT29^Scr^) cells (Fig. 2B). shRNA-induced SQOR knockdown led to 90-95% decrease in protein expression^16^ (Fig. S2A,B). From the sigmoidal dependence on sulfide concentration, an IC_50_ value of 30 ± 1 µM was estimated (Hill coefficient = 4.8) for HT29^Scr^ cells, and 6.0 ± 0.3 µM (Hill coefficient of 5.2) for HT29^SQOR^ ^KD^ cells (Fig. 2B). These data provide a quantitative measure of the protection afforded by SQOR against sulfide poisoning in HT29 cells.

**Figure 2.**
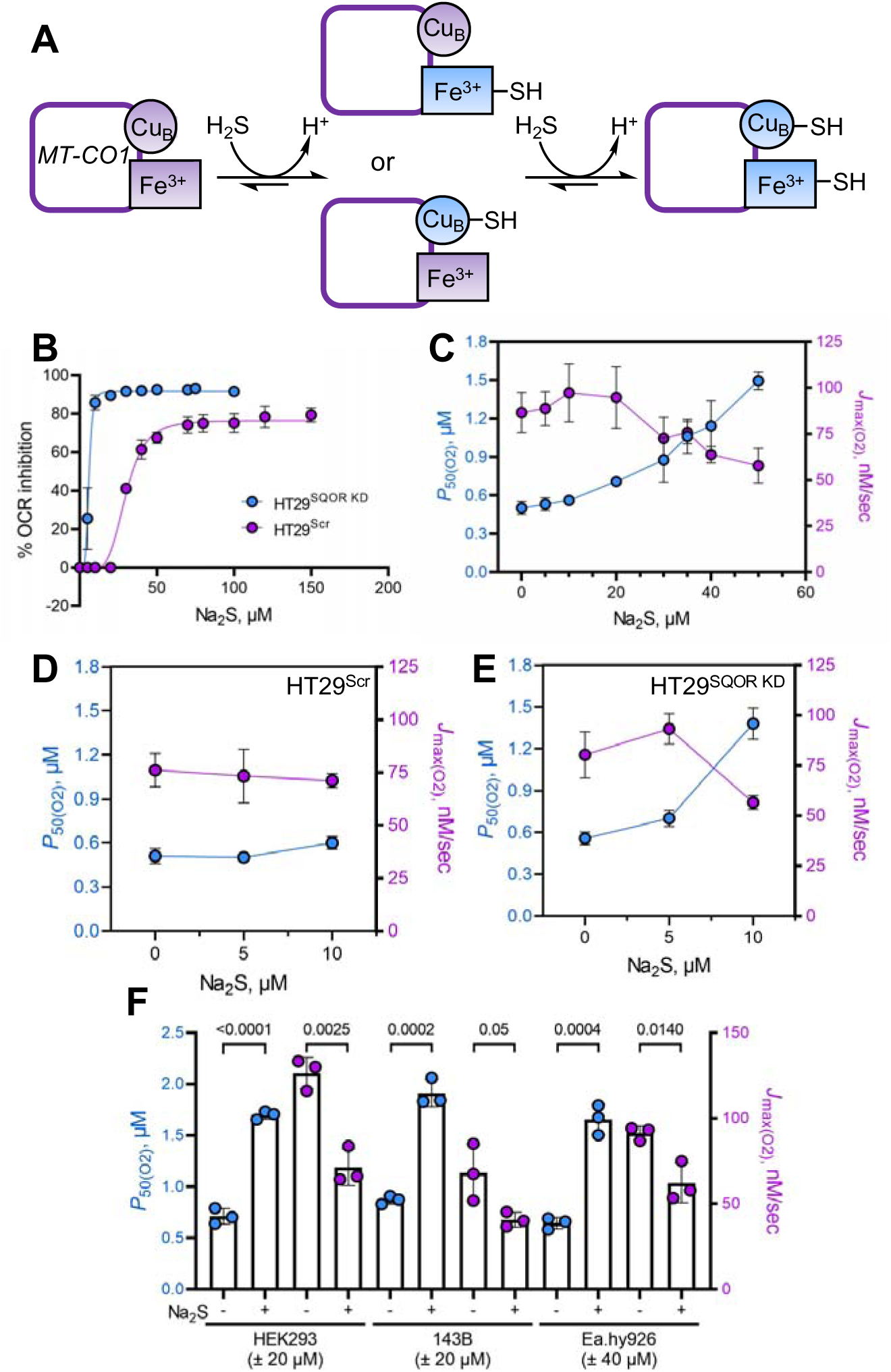
H_2_S increases P_50(O2)_ for complex IV. **A**. Postulated mechanism for sulfide binding to the metal sites in the MT-CO1 subunit of complex IV. **B**. Inhibition of OCR as a function of sulfide concentration in HT29^Scr^ (purple) and HT29^SQOR^ ^KD^ (blue) cells yielded IC_50_ estimates of 30 ± 1 and 6.0 ± 0.3 µM, respectively. Data represent the mean ± SD of 3-5 independent experiments. **C**. Dependence of *P*_50(O2)_ (blue) and *J*_max(O2)_ (purple) values on sulfide concentration in HT29 cells. Data represent the mean ± SD of at least 3 independent experiments. **D,E**. Na_2_S increases the *P*_50(O2)_ and decreases the *J*_max(O2)_ value in HT29^SQOR^ ^KD^ (at 10 µM) but not in HT29^Scr^ cells (n= 4). **F**. The *P*_50(O2)_ and *J*_max(O2)_ of HEK293 and 143B cells are significantly altered by 20 µM Na_2_S as in HT29 cells, while Ea.hy926 cells need 40 µM Na_2_S for similar modulation (n=3).

### H_2_S increases *P*_50(O2)_ for complex IV

Modulation of the O_2_ pressure at half maximal respiration, a measure of O_2_ affinity, was examined in HT29 cells. The *P*_50(O2)_ value increased from 0.50 ± 0.05 to 1.49 ± 0.07 µM while *J*_max(O2)_ decreased from 90 ± 10 to 58 ± 9 nM sec^-1^ at 37 °C, as sulfide concentrations increased from 0-50 µM (Fig. 2C). Technical issues precluded determination of *P*_50_ values at higher sulfide concentrations as the dissolved O_2_ was depleted during the recovery time, i.e. before cells could establish a new stationary OCR. Due to the greater sensitivity of HT29^SQOR^ ^KD^ cells, *P*_50(O2)_ modulation could only be evaluated at low Na_2_S (5-10 µM) concentrations (Fig. 2D,E). In contrast to HT29^Scr^ cells (Fig. 2D), a 1.3-fold increase in *P*_50(O2)_ (0.56 ± 0.05 versus 0.70 ± 0.06 µM) was seen with 5 µM Na_2_S, while the *J*_max(O2)_ value was unchanged (89 ± 9 versus 93 ± 8 nM sec^-1^). A 2.5-fold increase in *P*_50(O2)_ (1.4 ± 0.1 µM) was observed at 10 µM Na_2_S, which was accompanied by a 30% decrease in the *J*_max(O2)_ value (57 ± 3 nM sec^-1^) (Fig. 2E).

The ability of sulfide to modulate O_2_ metabolism was compared across cell lines (Fig. 2F). In HEK293 and 143B cells, 20 µM Na_2_S sulfide was sufficient to achieve a similar magnitude of *P*_50(O2)_ modulation as 40 µM Na_2_S in Ea.hy926 and HT29 cells. The *P*_50(O2)_ value increased 2.4-fold (0.71 ± 0.08 to 1.70 ± 0.04 µM) in HEK293 cells while *J*_max(O2)_ decreased from 126 ± 9 to 71 ± 11 nM sec^-1^. In 143B cells, *P*_50(O2)_ increased 2.2-fold (from 0.87 ± 0.04 to 1.91 ± 0.13 µM) while *J*_max(O2)_ decreased from 68 ± 17 to 41 ± 4 nM sec^-1^. In Ea.hy926 cells, *P*_50(O2)_ increased 2.5-fold (from 0.64 ± 0.06 to 1.7 ± 0.1 µM) and *J*_max(O2)_ decreased from 91 ± 4 to 60 ± 10 nM sec^-^1.

### A single acute H_2_S exposure elicits long-lived effects on O_2_ metabolism

We examined the durability of metabolic effects following a single acute exposure to Na_2_S (100 µM) after 4, 24 or 48 h (Fig. 3A,B). Perturbed OCR kinetics were clearly visible even 48 h later as evidenced by an increased sensitivity to subsequent Na_2_S exposure. A measurable increase in the *P*_50(O2)_ value was seen 24 h after a single exposure to Na_2_S, but the effect was dissipated at 48 h (Fig. 3C). Sulfide preconditioning decreased H_2_S consumption and thiosulfate production (Fig. 3D).

**Figure 3.**
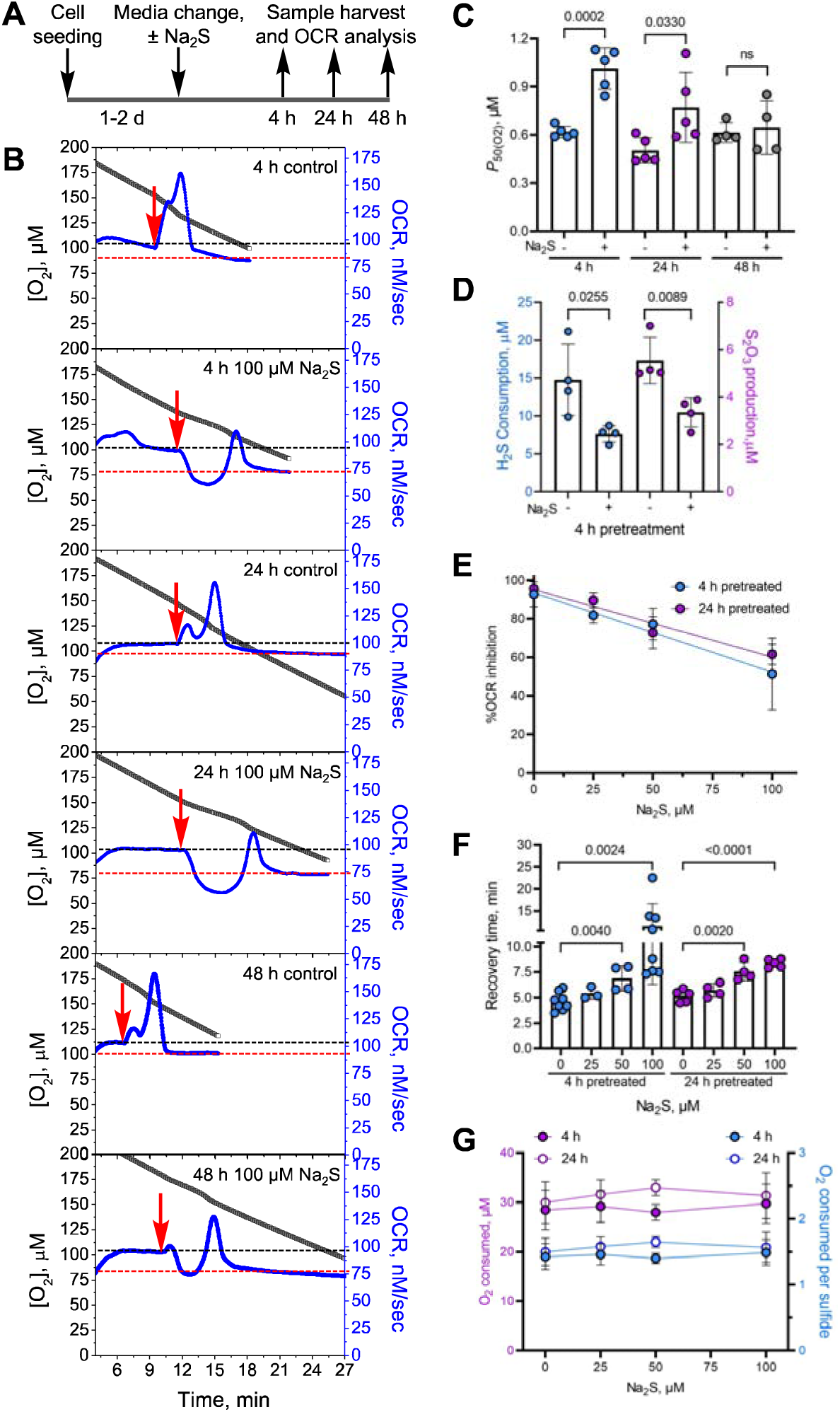
H_2_S memory is long-lived. **A**. Scheme showing the experimental setup used to test the durability of sulfide exposure. **B**. Representative OCR traces for cells ± 100 µM Na_2_S after 4, 24, and 48 h. While control cells rapidly oxidized 20 µM Na_2_S (red arrow), pretreated cells were inhibited up to 48 h. The black and red lines represent the OCR before and after recovery from 20 µM Na_2_S, respectively. **C**. Changes in cellular *P*_50(O2)_ values 4, 24 and 48 h after a single exposure to 100 µM Na_2_S. **D**. Pretreatment with 100 µM Na_2_S results in lower H_2_S clearance and thiosulfate production 4 h later. **E,F,G**. Sulfide (0-100 µM) pretreatment (4 or 24 h) resulted in a dose-dependent inhibition of OCR following Na_2_S re-exposure (20 µM) (E), increased time to recovery of a new stationary OCR (F), but had no impact on the total O_2_ consumed between sulfide and the recovery (G). The average total O_2_ consumed per sulfide was estimated to be 1.5 ± 0.1 across all sulfide pretreatment concentrations. Two-sample unpaired *t* test was performed for statistical analysis (C-G).

Next, the sensitivity of preconditioning to the sulfide dose was examined. Sulfide pretreatment (4 or 24 h) at concentrations as low as 25 µM H_2_S, increased sensitivity of OCR inhibition by a subsequent injection of sulfide (Fig 3E). The recovery time to a new stationary OCR, showed significant differences with ≥50 µM sulfide pre-treatment (Fig. 3F). Regardless of the duration of recovery, the control and sulfide pretreated cells consumed 30 ± 1 µM O_2_ before establishing a new stationary OCR, corresponding to an O_2_:sulfide ratio of ∼1.5 ± 0.1 (Fig. 3G).

Acute H_2_S exposure did not increase SQOR expression (Fig. S2A-C) or affect mitochondrial content as monitored by cardiolipin levels (Fig. S3A). Acute sulfide-exposure led to small but statistically significant decreases in MT-CO1 and MT-CO2 protein levels, which was correlated with slightly lower complex IV activity (Fig. S3B-D).

Cyanide also inhibits complex IV^29^. However, unlike sulfide, cyanide only coordinates to ferric or ferrous heme a_3_ but not to Cu_B_ in MT-CO1. The *K*_i_ for the *in vitro* inhibition of complex IV by cyanide is 0.2 µM, with *k*_on_ = 5 x 10^3^ M^-1^s^-1^ and *k*_off_ = 5 x 10^-4^ s^-1^ at 30°C (Fig. S4A)^28^. Surprisingly, cyanide pretreatment (500 µM, 4 h) enhanced sulfide-triggered OCR and recovery, following subsequent exposure to sulfide (Fig. S4B-D). These data suggest a specific mechanism for encoding memory by H_2_S that is not general to complex IV inhibition.

### Durability of H_2_S preconditioning correlates with SQOR expression levels

We examined whether an ∼10-fold difference in SQOR expression levels across five cell lines (Fig. S5A) are correlated with the durability of the preconditioning effect. The HEK293 cell line has very low SQOR expression and showed prolonged inhibition in response to Na_2_S over ∼50 min (Fig. S5B). Similarly, pretreatment of HT29^SQOR^ ^KD^ but not HT29^Scr^ control cells with sulfide for 4 or 24h led to sustained inhibition of OCR after a subsequent injection with a low (10 µM) Na_2_S (Fig. S2D-G). Panc-1 cells showed the highest SQOR expression and was the least sensitive to the preconditioning effect (Fig. S5C). However, HT29, LoVo and Ea.hy926 cells showed variable sensitivity to sulfide preconditioning although they had similar SQOR levels, suggesting that additional factors influence cellular H_2_S memory.

### H_2_S induces long-lived changes in mitochondrial function

Mitochondrial function analysis revealed pleiotropic changes associated with H_2_S memory (100 µM, 4 h) in HT29, HT29^Scr^ and HT29^SQOR^ ^KD^ cells (Fig. 4A and Fig. S6A). Basal, ATP-linked, and maximal respiration as well as the proton leak rate decreased, while non-mitochondrial respiration was unaffected, except in HT29^SQOR^ ^KD^ cells (Fig. 4B-G, Fig. S6B-G). Each of these parameters was more impacted in HT29^SQOR^ ^KD^ relative to HT29^Scr^ controls.

**Figure 4.**
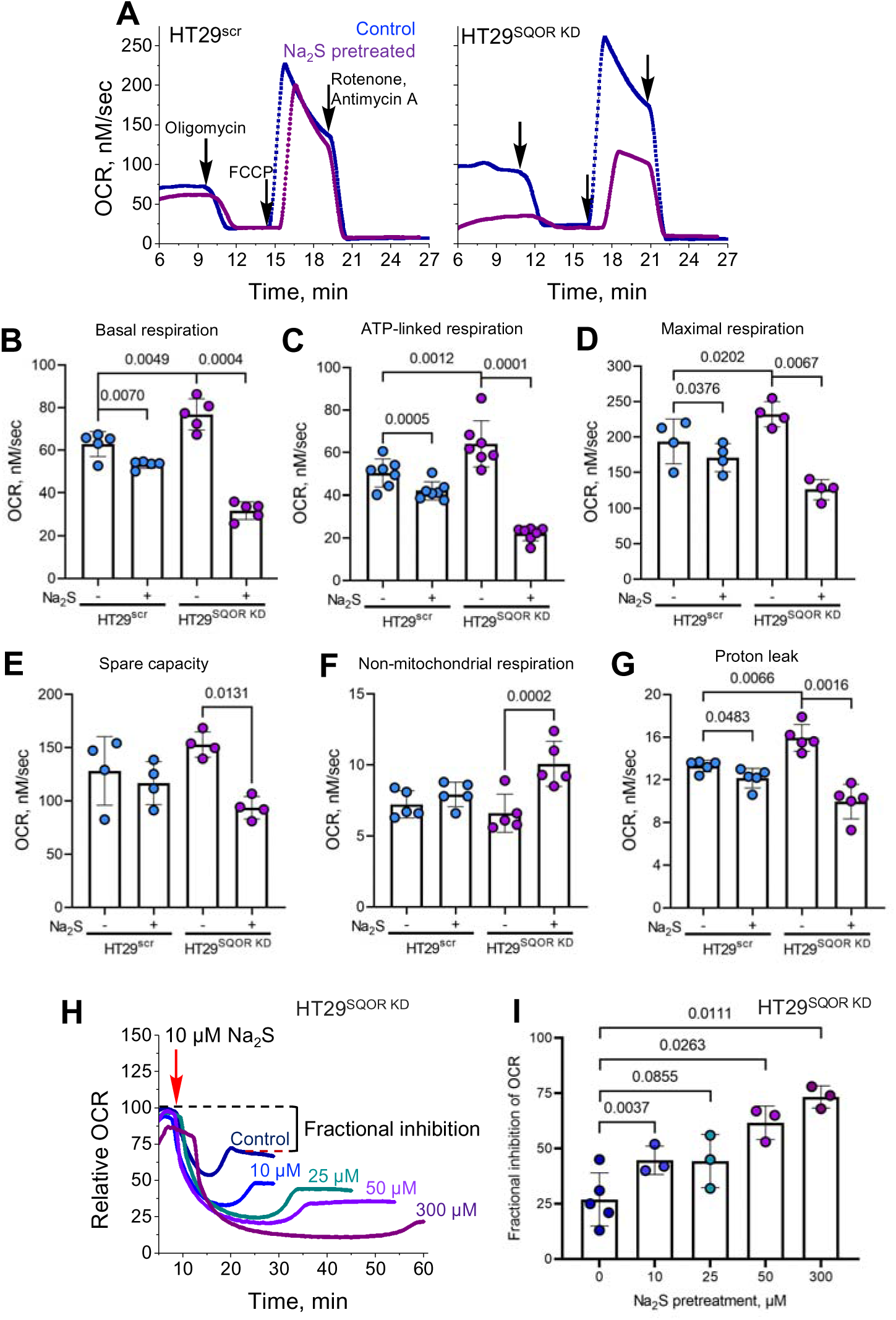
SQOR modulates impact of H_2_S on mitochondrial function. Representative traces of mitochondrial function profiles in HT29^scr^ (left panel) and HT29^SQOR^ ^KD^ (right panel) cells treated for 4 h ± 100 µM Na_2_S in response to oligomycin (125 nM), FCCP (125 nM), and rotenone and antimycin A (0.5 µM each). **B-G**. Quantitative analysis of the data in A reveals the effects of sulfide pretreatment on basal respiration (B), ATP-linked respiration (C), maximal respiration (D), spare capacity (E), non-mitochondrial respiration (F) and proton leak rate (G). **H, I**. Dose-dependent effects of Na_2_S pretreatment (10-300 µM, 24 h) on OCR (H) and the fraction of OCR inhibited (I) in HT29^SQOR^ ^KD^ cells following exposure to 10 µM Na_2_S (red arrow). Two-sample paired *t* test was performed for statistical analysis.

Prolonged and dose-dependent perturbations in mitochondrial bioenergetics were observed in HT29^SQOR^ ^KD^ cells after sulfide pretreatment (10-300 µM, 24 h). Enhanced sensitivity to respiratory inhibition was evident at concentrations as low as 10 µM Na_2_S, and a pronounced decrease in the stationary OCR, indicative of fractional inhibition was seen (Fig. 4H,I).

### H_2_S induces long-lived metabolic changes

Metabolomics analysis revealed long-lived changes in numerous metabolites (Fig. 5A), including some in the pentose phosphate and pyrimidine biosynthesis pathways (Fig. 5B,C), as well as an ∼50% decrease in serine levels (Fig. S7A).We have previously shown that sulfide induces an ∼2-fold increase in the glycolytic rate in HT29 cells to compensate for decreased ATP-linked respiration^22^. Remarkably, the dose-dependent increase in glucose consumption and lactate production persisted 24 h after an initial exposure to 25-300 µM sulfide (Fig. 5D,E). We have also shown that repeated exposure to sulfide increases incorporation of glutamine carbon to lipids via reductive carboxylation^16, 21^. Surprisingly, a single exposure to 100 or 300 µM Na_2_S was sufficient to increase [^14^C]-glutamine incorporation into lipids by 70 and 150%, respectively (Fig. 5F).

**Figure 5.**
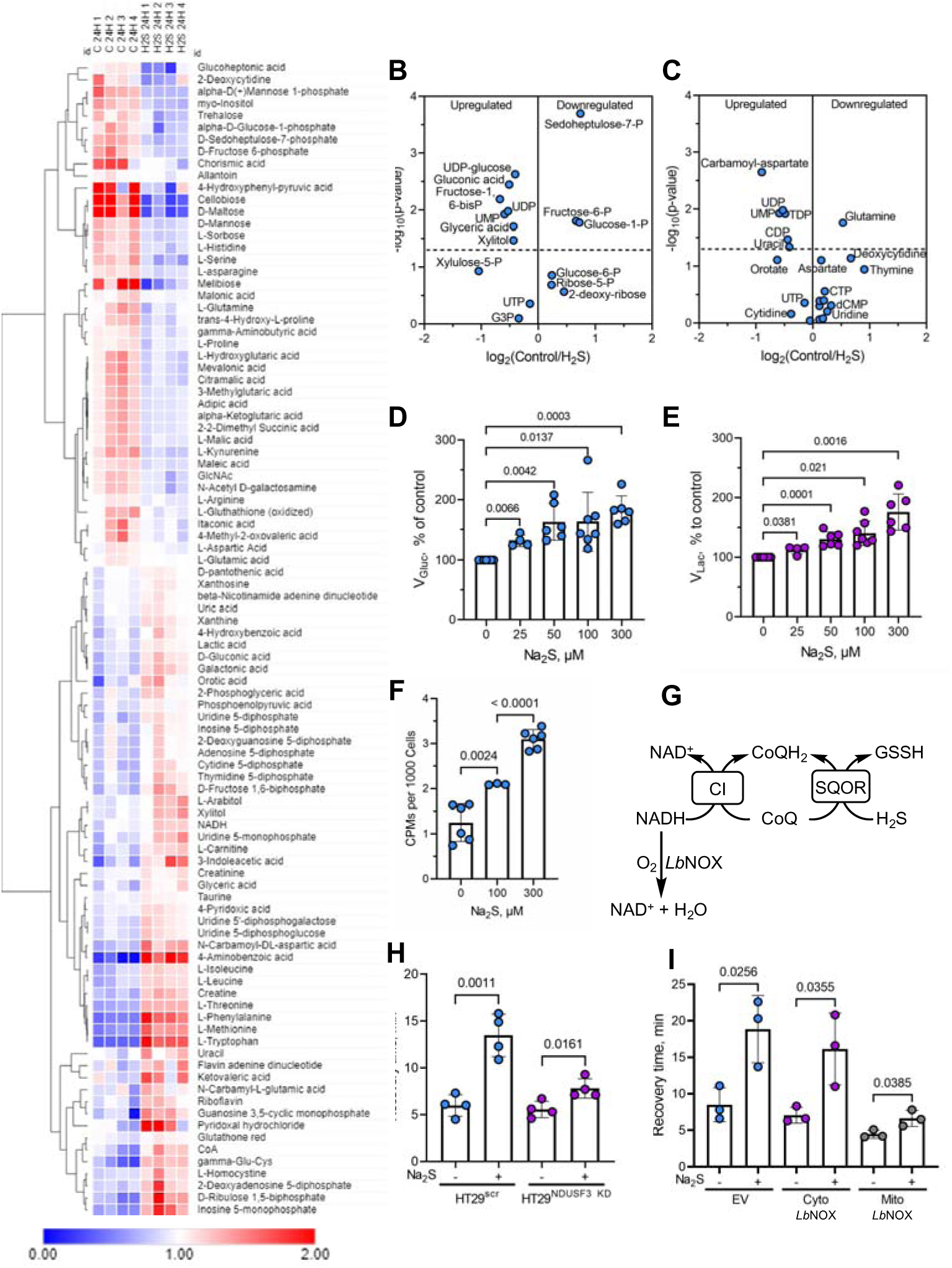
Sulfide preconditioning induces long-term metabolic changes implicating electron acceptor insufficiency. **A**. Metabolomics analysis in HT29 cells 24 h after ± 100 µM Na_2_S exposure. **B,C.** Changes in pentose phosphate (B) and pyrimidine biosynthesis (C) pathway metabolites are among the changes observed in the metabolomics profile. **D,E.** Pretreatment with sulfide (25-300 µM, 24 h) increased the relative rates of glucose consumption (D) and lactate production (E). Data are representative of at least 4 independent experiments. **F**. A single bolus exposure to sulfide (100 or 300 µM) increased [U^14^-C]-glutamine incorporation into lipids 13 h later (n*≥*3). **G**. Scheme showing how knockdown of the complex I subunit, NDUSF3, or mitochondrial *LbNOX* expression can impact CoQ availability. **H,I**. NDUSF3 knockdown (H) and expression of mitochondrial but not cytosolic *LbNOX* (I) decreased recovery time.

### Electron acceptor insufficiency contributes to H_2_S memory

A backup in the ETC is expected to cause an upstream reductive shift, resulting in electron acceptor insufficiency. We tested the hypothesis that CoQH_2_ recycling is limiting during the period that cells exhibit H_2_S memory. For this, electron transfer from complex I to CoQ was limited either by knockdown of the NDUFS3 subunit of complex I or via heterologous expression of *Lb*NOX, a water-generating NADH oxidase^30^, which can be targeted to the cytoplasm or mitochondria (Fig. 5G). In comparison to the HT29^Scr^ controls, H_2_S memory was substantially attenuated in HT29^NDUFS3^ ^KD^ cells as monitored by recovery time (Fig. 5H). Mitochondrial but not cytoplasmic *Lb*NOX expression also decreased recovery time compared to the empty vector control (Fig. 5I). Complex II reversal with fumarate, serving as an electron acceptor, can recycle CoQH_2_^26^ during acute H_2_S exposure (Fig. S7B). However, neither knockdown of the SDHA subunit of complex II nor provision of dimethylfumarate, a membrane-permeable fumarate derivative, alleviated H_2_S memory (Fig. S7C,D).

## DISCUSSION

The remarkable response of mice to moderate H_2_S exposure has the hallmarks of inducing hibernation-like behavior, including low core body temperature and depressed cardiac and metabolic function^6, 27, 31^. A study using a synthetic model of cytochrome c oxidase ascribed the molecular basis of these changes to reversible sulfide coordination at ferrous heme a_3_ in a manner that is competitive with O_2_^32^. While characterizing the short-term consequences of H_2_S exposure in a cell culture model in which systems like SQOR that interact with H_2_S are present, we discovered that H_2_S-induced metabolic perturbations endure over 4-48 h. The durability of this cellular memory, which is sensitive to H_2_S dose during pretreatment and the intrinsic capacity for sulfide oxidation by SQOR, was observed across all tested cell lines.

The persistence of cellular sulfide effects is surprising because H_2_S disappears rapidly under culture conditions (t^1^/_2_ ∼4 min in 6-cm plates, 37 °C)^25^. The durability of the preconditioning effects is also surprising since conserved stress response pathways exist to sense and alleviate acute reductive stress to preserve cellular integrity. For example, the redox status of conserved cysteines on FNIP1 (folliculin interacting protein 1) in myoblasts is sensed by the E3 ubiquitin ligase CUL2^FEM1B^, leading to FNIP degradation under reductive stress, with a consequent increase in mitochondrial activity^33^. Despite recent advances in our understanding of the acute effects of H_2_S on cellular metabolism^4, 5, 34^, the durability of the stress response to this volatile metabolite is not known.

The interaction of sulfide with the MT-CO1 subunit of complex IV is complex and poorly characterized^28^, and much less is known about its interaction with the di-copper-containing MT-CO2 subunit. Sulfide reacts with fully oxidized MT-CO1 to give ferric heme a_3_ with a sulfide ligand and cuprous Cu_B_^35^. Sulfide oxidation can occur from this mixed valence state of MT-CO1 although the product, which could be HSSH, polysulfide, S^0^ or S_8_, remains to be characterized Fig. 6A). Sulfide binding and oxidation to catenated sulfur species has been described for other hemeproteins^36–39^.

**Figure 6.**
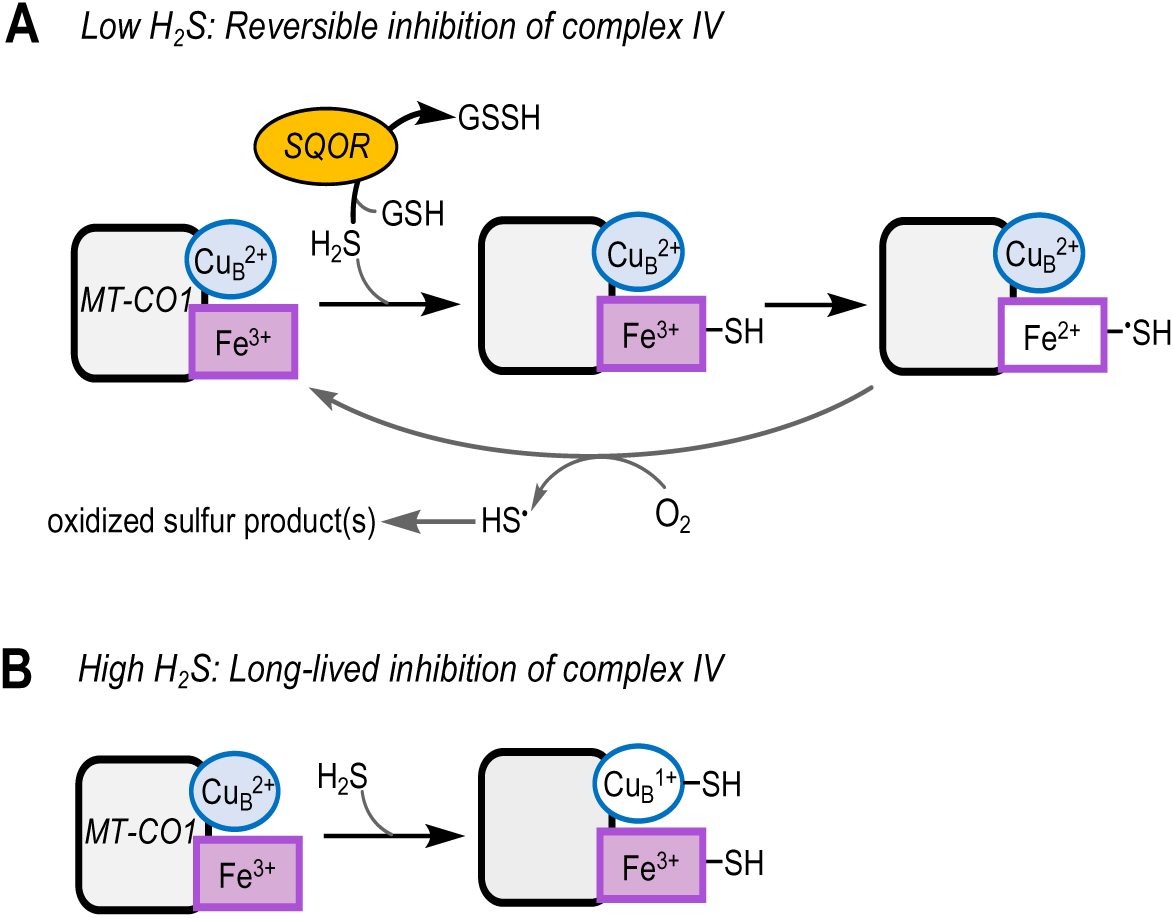
Summary of H_2_S interactions with MT-CO1 at low and high H_2_S concentrations. **A.** At low sulfide concentrations, H_2_S is efficiently cleared by SQOR and its interaction with MT-CO1 leads to weak and reversible inhibition that is competitive with respect to O_2_. **B**. When SQOR capacity is exceeded and H_2_S levels rise, coordination to the Cu_B_ site in MT-CO1 leads to long-lived inhibition of complex IV.

In a cellular milieu, the interaction of H_2_S with complex IV is modulated by SQOR^19^. We posit that at low concentrations, SQOR limits H_2_S exposure, leading to reversible complex IV inhibition via formation of ferrous heme a_3_ (Fig. 6A). When the capacity to clear sulfide is limiting or compromised, as in SQOR deficiency, H_2_S concentrations rise and MT-CO1 containing ferric heme a_3_ and cuprous Cu_B_-sulfide accumulates (Fig. 6B). Spectroscopic analysis of sulfide-treated MT-CO1 reveals that the Cu_B_ site is resistant to air oxidation, even when the heme centers have been re-oxidized, which has been interpreted as evidence for an increase in the redox potential of cuprous Cu_B_ via direct sulfide coordination^40^. Long-lived fractional inhibition of complex IV following H_2_S preconditioning, would explain lower basal OCR, ATP-linked respiration, and spare respiratory capacity as well as higher *P*_50(O2)_ values, (Figs. 3C and 4, Fig. S6). Sustained fractional inhibition of complex IV is also supported by the statistically significant decrease in its activity (Fig. S3D), which could be an underestimate since ascorbate/TMPD could promote loss of the sulfide ligand. While the effect of sulfide-coordination on the stability of MT-CO1 or MT-CO2 is unknown, we note small but statistically significant decreases in protein levels just 4 h after an acute sulfide exposure (Fig. S3B,C). Our model also explains why cyanide, which interacts with the heme but not the copper site, fails to elicit comparable long-lived changes (Fig. S4).

The cellular response to H_2_S ranges from stimulation of O_2_ consumption at low, to inhibition at high concentrations, while complex kinetics are observed at intermediate concentrations, which reflect the dual ETC substrate and inhibitor dynamic of this metabolite. Regardless of the initial response, cells appear to return to basal OCR, superficially suggesting reversibility. However, a closer inspection reveals fractionally lower stationary OCR values after successive exposure to H_2_S, which is more evident at the higher concentrations (Fig. 1B) and marked in HT29^SQOR^ ^KD^ cells (Fig. S2D,F). Persistent dampening of the ETC flux after sulfide injection is more readily seen in sulfide-pretreated cells (Fig. 3B) and particularly, in HT29^SQOR^ ^KD^ cells (Fig. 4H,I), raising questions as to the underlying mechanism.

The observed increase in *P*_50(O2)_ with a concomitant decrease in *J*_max(O2)_ (Fig. 2C), is inconsistent with noncompetitive inhibition by sulfide with respect to O_2_, as reported in an *in vitro* steady-state kinetic analysis^29^. It has been argued that the slow off-rate for sulfide likely distorted the kinetic pattern, decreasing *V*_max_ while having a minimal effect on the *K*_M(O2)_ due to an increasing fraction of the enzyme being inactive with increasing sulfide concentration^28^. We posit that the ability of sulfide to increase *P*_50(O2)_ for complex IV explains the protective effects of sulfide pre-conditioning on lethal hypoxia^27^ as well as limiting reperfusion injury following ischemia^8^. Decreased ETC flux is expected to protect against oxidative injury.

SQOR releases two electrons per catalytic cycle, generating one equivalent of COQH_2_ that leads to the reduction of 0.5 O_2_ per mole of sulfide oxidized. Since glutathione persulfide, the other product of the SQOR reaction, is further oxidized to sulfite, consuming one mole of O_2_, the predicted stoichiometry of O_2_ consumed:sulfide is 1.5. However, if H_2_S also serves as an alternate sulfane sulfur acceptor, generating HSSH^20^, the expected O_2_:sulfide (1.5:2.0) stoichiometry is 0.75. The experimentally determined ratio was 1.0 ± 0.1 in rotenone-treated cells (Fig. 1C), and 1.5 ± 0.2 in sulfide-preconditioned cells (in the absence of rotenone) (Fig. 3G). Both values are higher than the 0.74-0.89 stoichiometry reported previously^41^. Loss through volatilization of H_2_S is unlikely to be a significant factor in the closed respirometry chamber used in our study. Furthermore, a larger fractional deviation from the expected stoichiometry would be expected due to H_2_S loss at lower sulfide concentrations, which was not observed. Deviation from the 1.5 stoichiometry in rotenone-treated cells could be explained by: (i) incomplete coupling between the SQOR and ETHE1-catalyzed reactions, or between SQOR and complex III activity (Fig 1A), or (ii) the dual use of H_2_S (*k*_cat_/*K*_M_ = 3.7 x 10^5^ M^-1^ s^-1^) and GSH (*k*_cat_/*K*_M_ = 1.6 x 10^4^ M^-1^ s^-1^)^20^ as sulfur acceptors during an acute H_2_S exposure, or (iii) diversion of CoQH_2_ to complex II functioning in reverse (Fig. S7B)^26^.

We have previously reported the profound effect of SQOR in protecting against sulfide inhibition of the ETC^19^ and characterized it quantitatively in this study. SQOR deficiency decreased the IC_50_ for H_2_S from 30 ± 1 to 6.0 ± 0.3 µM in HT29 cells and rendered the *P*_50(O2)_ value sensitive to low H_2_S concentrations (*≤* 10 µM) in SQOR knockdown but not in control cells (Fig. 2B,D,E). Since the knockdown efficiency was ∼90-95%, our study underestimates the full extent of ETC modulation by SQOR in these cells. Patients with SQOR deficiency present with Leigh disease and limited tissue analysis shows markedly lower complex IV activity^42^. The protein levels of only the nuclear encoded subunits were investigated in this study and found to be normal, leading to the conclusion that complex IV activity, but not its assembly, was affected by SQOR deficiency. Ethylmalonic encephalopathy, another inborn error of metabolism characterized by elevated H_2_S, results from deficiency of ETHE1, the second enzyme in the sulfide oxidation pathway (Fig. 1A)^43^. The disease is characterized by severe deficiency of complex IV activity particularly in brain and muscle, and decreased MT-CO1 and MT-CO2 protein levels^44^. These studies provide additional support of our finding that acute H_2_S exposure has long-lived effects on complex IV, which are exacerbated in the absence of a functional sulfide oxidation pathway.

In summary, our study provides a quantitative analysis of ETC regulation by H_2_S via decreased O_2_ affinity, sustained complex IV inhibition, and partial destabilization of the metal-containing MT-CO1 and MT-CO2 subunits. SQOR, which is a key determinant of cellular H_2_S levels^14^, could represent both an “on” and an “off” switch for sulfide-dependent ETC regulation^34^. The prolonged effects on oxidative metabolism provide an opportunity for extending the time window in which the potential therapeutic effects of H_2_S for attenuating injury or prolonging platelet shelf life might be exploited.

## METHODS

### Materials

Na_2_S, nonahydrate (99.99% purity, (431648), rotenone (R8775), protease inhibitor cocktail for mammalian tissue extract (P8340), puromycin (P8833), dimethylfumarate (242926), doxycycline (D3447), hexane (110543), RIPA lysis buffer (R0278), dimethyl sulfoxide (D2650), oligomycin A (75351), antimycin A (A8674), and ascorbate (PHR1279-1G) were from Sigma. Dulbecco’s modified Eagle’s medium (DMEM) (with 4.5 g/L glucose, 584 mg/L glutamine, and 110 mg/L sodium pyruvate, (11995-065)), RPMI 1640 with glutamine (11875-093), fetal bovine serum (FBS, 10437-028), penicillin/streptomycin mixture (15140-122), 0.05% (w/v) trypsin-EDTA (25300-054), phenol red free 0.5% trypsin EDTA (15400054), PBS (10010-023), and Dulbecco’s phosphate-buffered saline medium (DPBS, 14040-133) were from Gibco. Geneticin (10131-035) was from Life Technologies. N,N,N’,N’-tetramethyl-*p*-phenylenediamine dihydrochloride (TPMD, T01511G), methanol (A452-4), molecular grade isopropanol (BP2618-500), molecular grade HEPES (7365-45-9), and Tween20 (BP337500) were from Fisher. [U-^14^C]-glutamine (281.0 mCi/mmol) was from PerkinElmer. KCl (7447-40-7) was from Acros Organics and Nonidet P40 substitute (74385) was from Fluka BioChemika. Nonyl acridine orange (NAO, A1372) was purchased from Invitrogen. The D-glucose detection kit (K-GluHK-110A) was from Megazyme, Carbonyl cyanide-p-trifluoromethoxyphenylhydrazone (FCCP, 15218) and the L-lactate assay kit (700510) were from Cayman Chemical. 10% Pre-cast tris-glycine gels (4561033), PVDF membranes (162-0177), thick blot filter paper (1703932), and clarity ECL substrate (1705062) were purchased from Bio-Rad. Anti-SQOR antibody (17256-1-AP) was from Proteintech, anti-MTC-O1 and anti-MT-CO2 antibodies (ab14705 and ab110258, respectively) were from Abcam. Anti-rabbit horseradish peroxidase-linked IgG and anti-mouse IgG, horseradish peroxidase linked antibodies (NA944V and NA931, respectively) were from GE Healthcare. Polystyrene round-bottom tubes (5 mL) with cell-strainer caps used for FACS analysis was from BD Biosciences (720035).

### Cell culture

HT29 and Panc-1 cells were maintained in RPMI 1640 medium. HEK293, RKO, LoVo, and Ea.hy926 cells were maintained in DMEM medium. Both RPMI and DMEM media were supplemented with 10% FBS along with 100 units/mL penicillin and 100 µg/mL streptomycin. All cells were maintained at 37°C with ambient O_2_ and 5% CO_2_. A separate incubator was used for cells exposed to Na_2_S to avoid cross contamination of control samples with the volatile metabolite. HT29^scr^, HT29^SQOR^ ^KD^, HT29^NDUSF3^ ^KD^, and HT29^SDHA^ ^KD^ cell lines were maintained in the same medium as the parent HT29 cells but with 1 µg/mL puromycin. Puromycin was removed from the culture medium for 2-3 day experiments to avoid potential off target effects of the additional antibiotic. HT29 cells expressing the cytosolic or mitochondrial *Lb*NOX were cultured as described previously^24^ with geneticin antibiotic (300 µg/mL) added for selection and 300 ng/mL doxycycline used for induction 24 h before the start of the experiment.

### Oxygen consumption rate (OCR) measurements

All OCR measurements were performed using a respirometer (Oroboros Instruments Corp) at 37 °C with a stirring rate of 750 rpm. Cells were harvested from 10 cm plates (at ∼70-90% confluency) washed with 1 x 8 mL PBS prior to digestion with 1 mL trypsin (0.05%) at 37°C for 5-10 min. Cells were collected in 7 mL of the cell culture medium and centrifuged at 1600 x *g* for 5 min, and pellets were resuspended in 1 mL modified PBS (MPBS) or DPBS + 5 mM glucose + 20 mM HEPES, pH 7.4. Cell suspensions were transferred to pre-weighed tubes, centrifuged at 1600 x *g* for 3 min and the supernatant was carefully aspirated with a 2 µL tip fixed to a vacuum line. The wet weight of the pellet was determined to prepare 5 % (w/v) cell suspensions in MPBS, which were kept on ice and used for dilution to 0.75% or 1% (w/v) suspensions for OCR experiments. Experiments in which 0.75% (w/v) cell suspensions were used included those performed in the presence of rotenone (before Na_2_S treatment) or with *Lb*NOX expressing cells, and in experiments in which mitochondrial function was assessed.

Analysis of OCR traces was performed using DatLab v6 (Oroboros Instruments, Austria) and replotted in Origin 7.0. Recovery time following Na_2_S injection is defined as the time taken by cells to return to a new stationary basal OCR. Sulfide-dependent (% OCR) inhibition refers to the maximal drop in OCR following sulfide injection and is denoted as a percent of the starting basal OCR (Figs. 2B and 3E). Fractional inhibition of OCR refers to the change in basal OCR following recovery from sulfide-induced inhibition and is expressed as a percent difference between the basal OCR at the start of the experiment and the new stationary basal OCR. When calculating the ratio of O_2_ consumed per sulfide added, the contribution of non-mitochondrial respiration, which was determined to be 10% of the basal OCR in HT29 cells, was subtracted from the total O_2_ consumed before dividing by the concentration of sulfide added (5-35 µM). The ΔOCR_max_ represents the maximal change in OCR in response to 5-35 µM Na_2_S in rotenone treated cells.

### Mitochondrial function analysis

The two chambers in the Oroboros instrument were filled with control versus 100 µM Na_2_S pretreated (4 h) HT29, HT29^SQOR^ ^KD^ or HT29^scr^ cell suspensions (0.75%, 2 mL total volume). Once basal OCR stabilized, 1 µL of 250 µM oligomycin (125 nM final concentration) was injected into each chamber to inhibit ATP-linked respiration. After then new OCR stabilized (∼3 min), 1 µL of 250 µM FCCP (125 nM) was added to elicit maximal respiration, followed 2-3 min later by 0.5 µL of a combined stock of 2 mM rotenone + 2 mM antimycin A (0.5 µM each, final concentration). Working stocks of oligomycin (250 µM), FCCP (250 µM) and rotenone + antimycin A (2 mM each) were prepared in cell culture grade DMSO, aliquoted, stored at -20 °C and thawed for single use.

### Complex IV activity measurements using the TMPD assay

Complex IV activity was measured in 1% (w/v) cell suspensions of control and Na_2_S pretreated (100 µM, 4 h) cells in the Oroboros instrument as described previously^45^. Briefly, once basal OCR stabilized, complexes I and III were blocked with rotenone and antimycin A (0.5 µM each final concentration). After a new stable OCR was established, FCCP (1 µM) was added, followed after a few minutes by ascorbate (200 µM) and then, N,N,N’,N’-tetramethyl-p-phenylenediamine dihydrochloride (TMPD, 80 µM). TMPD leads to a spike in the OCR, corresponding to complex IV activity, and was recorded over at least 2 min. The ascorbate stock solution (200 mM) was prepared by diluting a freshly prepared 800 mM stock solution whose pH was adjusted to ≈6 with 5 N HCl.

### *P*_50(O2)_ and *J*_max(O2)_ analysis

Cell suspensions (1% (w/v)) were prepared as described above for all other OCR experiments. Temperature and stirrer settings were as noted (37 °C and 750 rpm). Cells were pretreated either with100 µM Na_2_S (4, 24 or 48 h) or Na_2_S (5-50 µM) was injected into the Oroboros chamber after stabilization of basal OCR (typically, 10-15 min). O_2_ concentration was recorded at 0.5 sec time intervals until O_2_ in the chamber was depleted and for an additional 5 min to determine the flux at zero O_2_ for background correction. DatLab v2 (Oroboros Instruments, Austria) was used to estimate the *P*_50(O2)_ (or apparent *K*_M(O2)_) and *J*_max(O2)_ (or maximum flux) values as described^46, 47^. Briefly, instrumental background parameters (zero intercept, a°, and slope, b°, of the background O_2_ flux as a function of O_2_ concentration) and time constants (*τ*, time delay due to diffusion of O_2_ through sensor membrane) for each chamber were determined experimentally according to the manufacturer’s instructions in the O2k-Manual. Time constants were determined via stirrer test and DatLab v2 macro TIMECONS, and O_2_ traces were recorded for 100% air calibration in MPBS at the start of each day when experiments were run. DatLab v2 macro P50 was used to estimate *P*_50(O2)_ and *J*_max(O2)_ values. Volume-specific OCR (oxygen flux, *J*_O2_) was calculated as the negative slope of the recorded oxygen concentration. After interpolation of zero flux, the macro iteratively fits the data to a hyperbolic equation given by: J_O2_ = (J_max_ × P_O2_)/(P_50_ + P_O2_)

where *J*_O2_ = volume-specific OCR and *P*_O2_ = [O_2_] in kPa. Oxygen partial pressure used for the fit was 0-1.1 kPa and oxygen solubility was 10.56 µM/kPa.

### IC_50_ for H_2_S

The IC_50_ for H_2_S was determined with 1% (w/v) HT29^SQOR^ ^KD^ or HT29^scr^ cell suspensions in MPBS in the Oroboros instrument. After the basal OCR stabilized at 37 °C, Na_2_S (30-150 µM for HT29^scr^ and 5-100 µM for HT29^SQOR^ ^KD^) was injected from a freshly prepared 10 mM stock solution. Sulfide-dependent % OCR inhibition was estimated by subtracting the lowest recorded OCR value from the basal OCR value, and expressed as a percentage of the starting basal OCR. From the dependence of % OCR inhibition on H_2_S concentration, the IC_50_ was obtained by fitting the data to a 4-parameter sigmoidal equation given by:

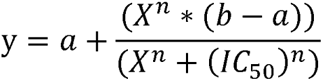

where “y” represents ΔOCR, “a” is the lowest value of ΔOCR, “b” is the highest value of ΔOCR and “n” is the Hill coefficient.

### Western blot analysis

*SQOR*. HT29^SQOR^ ^KD^ and HT29^scr^ were seeded at 500,000 cells per well in 6-well plates and cultured for 24-48 h before changing to fresh RPMI medium ± 100 µM Na_2_S for 4, 24 or 48 h. Cells that would be harvested after 4 h were seeded reach ∼70% confluency while cells that would be harvested after 24-48 h were seeded to reach ∼50% confluency before sulfide treatment. Cells were washed with 2 x 2 mL PBS/well followed by addition of 0.5 mL 0.05% trypsin/well for 5 min at 37 °C and harvested in 1 mL RPMI medium. Cell suspensions were centrifuged at 1600 x *g* for 5 min and the pellet was washed once with 1 mL ice-cold PBS and resuspended in 200 µL RIPA lysis buffer + protease inhibitor and saved at -80°C until lysis. For SQOR detection in different cell lines, a similar procedure was used but each cell line was cultured in 10 cm plates and grown to confluency. Cells were washed with 8 mL PBS/plate followed by addition of 1 mL 0.05% trypsin for 5-10 min and harvested in 7 mL RPMI. Cells were centrifuged at 1600 x *g* for 5 min, the pellet was washed with 1 x 1 mL ice cold PBS and resuspended in 300 µL non-denaturing lysis buffer with protease inhibitor (0.5% (v/v), Nonidet P40 substitute, 25 mM KCl, 20 mM HEPES, pH 7.4) with the exception of Ea.hy926 cells, which were suspended in 100 µL of lysis buffer, due to the smaller pellet size.

*MT-CO1 and MT-CO2*. Cells (2 million per 6 cm dish) were grown for 36 h before changing to fresh RPMI medium ± 100 µM Na_2_S. After 4 h, the cells were harvested as described above and the pellet was resuspended in 400 µL non-denaturing Nonidet P40 lysis buffer with protease inhibitor.

#### Development of western blots

Frozen cell pellets were lysed by three freeze-thaw cycles and centrifuged at 13,000 x *g* for 10 min. Protein content in the supernatant was determined using the Bradford reagent (Bio-Rad). Cell lysates (50 µg) were loaded on precast Bio-Rad 10% tris-glycine gels and separated by SDS-PAGE, transferred to PVDF membranes, blocked for 1 h with 5% milk in Tris buffered saline with 0.3% Tween 20 (TBST). Membranes were soaked overnight in TBST, 5% (w/v) milk containing diluted SQOR (1:5000) or MTCO1 or MTCO2 (1:1000) antibodies then washed quickly with 2 x 5 mL TBST followed by 5 x10 min washes with 10-15 mL TBST. Then, the membranes were exposed for 2 h to the secondary antibody (horseradish peroxidase-linked anti-rabbit (for anti-SQOR) or anti-mouse IgG (for MTCO-1 and MTCO-2) used at a 1:10,000 dilution in TBST, 5% milk. The membranes were washed quickly with 2 x 5 mL TBST followed by 5 x 10 min 10-15 mL TBST washes, before rinsing quickly with 2x10 mL TBS before treating with clarity ECL substrate (Bio-Rad). Signals were detected using a Bio-Rad ChemiDoc Imaging System. All images were exported as 16-bit TIF files and imported into Fiji^48^ for semi-quantitative analysis, which was performed by drawing equal sized rectangles over each band to estimate its mean pixel intensity. The background was subtracted from each band.

### Cardiolipin analysis

Mitochondrial content was estimated by staining for cardiolipin with nonyl acridine orange (NAO) as described previously^19^. Briefly, HT29 cells were seeded at 700,000 cells/well in 6 well plates and the medium was changed after 24 h before ± 100 µM Na_2_S treatment. After 24 h, cells were washed once with and then replaced by phenol red free RPMI and treated ± 100 nM NAO for 30 min. Cells were then washed twice with 1 mL ice-cold PBS, treated with 0.5 mL phenol red free trypsin EDTA (0.05%) for 5 min, and harvested with 1 mL phenol red free RPMI. Cells were centrifuged at 700 x *g* or 5 min and the pellet was resuspended in 750 µL ice-cold PBS by pipetting and then filtered using 5 mL BD round bottom falcon tubes with cell-strainer caps. FACS analysis was performed on the Bio-Rad Ze5 multi-laser, high speed cell analyzer operated with the Everest software package at the University of Michigan Flow Cytometry Core Facility. Data were analyzed using FlowJo (v10.8.1), for median fluorescence (488 excitation 525/50 emission). The median unstained fluorescence was subtracted from the value for the stained sample.

### Metabolite analyses

Changes in glucose consumption and lactate production kinetics were measured in 5% (w/v) cell suspensions 24 h after a single bolus treatment with 25-300 µM Na_2_S exactly as described previously^25^. Incorporation of [U-^14^C]-glutamine into lipids was measured 13 h after a single bolus treatment with 100 or 300 µM Na_2_S as described^24^. H_2_S and thiosulfate concentrations in the conditioned medium were monitored 4 h after a single bolus treatment with 100 µM Na_2_S following derivatization with monobromobimane as previously described^26^. Metabolomics analysis was performed on HT29 cells treated ± 100 µM Na_2_S for 24 h, as described previously^19^.

### Statistics

Statistical analysis for the indicated pairwise comparisons of *P*_50(O2)_, *J*_max(O2)_, and protein abundance was performed using a two-sample unpaired t-test. For complex IV activity, parameters of mitochondrial function, glucose consumption, lactate production, and enhanced radioactive incorporation into lipids, paired t-tests were employed. Any other method used for statistical analysis is mentioned in the respective figure legend.

## ACKNOWLEDGEMENTS

This work was supported in part by grants from the National Institutes of Health (GM130183 to RB, R01CA248160 to CAL and F32GM140694 to DH) and the Michigan Life Sciences Fellow Program (to JD).

## DATA AND MATERIALS AVAILABILITY

All data are available in the manuscript or supplementary materials.

## COMPETING INTEREST STATEMENT

C.A.L. has received consulting fees from Astellas Pharmaceuticals, Odyssey Therapeutics, and T-Knife Therapeutics, and is an inventor on patents pertaining to Kras regulated metabolic pathways, redox control pathways in pancreatic cancer, and targeting the GOT1-pathway as a therapeutic approach (US Patent No: 2015126580-A1, 05/07/2015; US Patent No: 20190136238, 05/09/2019; International Patent No: WO2013177426-A2, 04/23/2015).

## REFERENCES

1. Morrison, M.L. et al. Surviving blood loss using hydrogen sulfide. J Trauma 65, 183–188 (2008).

2. Yenari, M.A. & Han, H.S. Neuroprotective mechanisms of hypothermia in brain ischaemia. Nat Rev Neurosci 13, 267–278 (2012).

3. Abe, K. & Kimura, H. The possible role of hydrogen sulfide as an endogenous neuromodulator. J Neurosci 16, 1066–1071 (1996).

4. Hanna, D., Kumar, R. & Banerjee, R. A Metabolic Paradigm for Hydrogen Sulfide Signaling via Electron Transport Chain Plasticity. Antioxid Redox Signal 38, 57–67 (2023).

5. Kumar, R. & Banerjee, R. Regulation of the redox metabolome and thiol proteome by hydrogen sulfide. Crit Rev Biochem Mol Biol 56, 221–235 (2021).

6. Blackstone, E., Morrison, M. & Roth, M.B. H_2_S induces a suspended animation-like state in mice. Science 308, 518 (2005).

7. Filipovic, M.R., Zivanovic, J., Alvarez, B. & Banerjee, R. Chemical Biology of H_2_S Signaling through Persulfidation. Chem Rev 118, 1253–1337 (2018).

8. Elrod, J.W. et al. Hydrogen sulfide attenuates myocardial ischemia-reperfusion injury by preservation of mitochondrial function. Proc Natl Acad Sci U S A 104, 15560–15565 (2007).

9. Marutani, E. et al. Sulfide catabolism ameliorates hypoxic brain injury. Nat Commun 12, 3108 (2021).

10. Linden, D.R. Hydrogen sulfide signaling in the gastrointestinal tract. Antioxid Redox Signal 20, 818–830 (2014).

11. Li, Z. et al. Mitochondrial H(2)S Regulates BCAA Catabolism in Heart Failure. Circ Res 131, 222–235 (2022).

12. Furne, J., Saeed, A. & Levitt, M.D. Whole tissue hydrogen sulfide concentrations are orders of magnitude lower than presently accepted values. Am J Physiol Regul Integr Comp Physiol 295, R1479–1485 (2008).

13. Levitt, M.D., Abdel-Rehim, M.S. & Furne, J. Free and Acid-labile Hydrogen Sulfide Concentrations in Mouse Tissues: Anomalously High Free Hydrogen Sulfide in Aortic Tissue. Antioxid Redox Signal 15, 373–378 (2011).

14. Vitvitsky, V., Kabil, O. & Banerjee, R. High turnover rates for hydrogen sulfide allow for rapid regulation of its tissue concentrations. Antioxid Red Signal 17, 22–31 (2012).

15. Deplancke, B. et al. Gastrointestinal and microbial responses to sulfate-supplemented drinking water in mice. Exp Biol Med (Maywood) 228, 424–433 (2003).

16. Macfarlane, G.T., Gibson, G.R. & Cummings, J.H. Comparison of fermentation reactions in different regions of the human colon. J Appl Bacteriol 72, 57–64 (1992).

17. Landry, A.P., Ballou, D.P. & Banerjee, R. Hydrogen sulfide oxidation by sulfide quinone oxidoreductase. Chembiochem 22, 949–960 (2021).

18. Landry, A.P., Roman, J. & Banerjee, R. Structural perspectives on H_2_S homeostasis. Curr Opin Struct Biol 71, 27–35 (2021).

19. Libiad, M. et al. Hydrogen sulfide perturbs mitochondrial bioenergetics and triggers metabolic reprogramming in colon cells. J Biol Chem 294, 12077–12090 (2019).

20. Landry, A.P., Ballou, D.P. & Banerjee, R. H_2_S oxidation by nanodisc-embedded human sulfide quinone oxidoreductase. J Biol Chem 292, 11641–11649 (2017).

21. Landry, A.P. et al. A catalytic trisulfide in human sulfide quinone oxidoreductase catalyzes coenzyme A persulfide synthesis and inhibits butyrate oxidation. Cell Chem Biol 26, 1515–1525 e1514 (2019).

22. Goubern, M., Andriamihaja, M., Nubel, T., Blachier, F. & Bouillaud, F. Sulfide, the first inorganic substrate for human cells. FASEB J 21, 1699–1706 (2007).

23. Nicholls, P. & Kim, J.K. Sulphide as an inhibitor and electron donor for the cytochrome c oxidase system. Can J Biochem 60, 613–623 (1982).

24. Carballal, S. et al. Hydrogen sulfide stimulates lipid biogenesis from glutamine that is dependent on the mitochondrial NAD(P)H pool. J Biol Chem, 100950 (2021).

25. Vitvitsky, V. et al. The mitochondrial NADH pool is involved in hydrogen sulfide signaling and stimulation of aerobic glycolysis. J Biol Chem, 100736 (2021).

26. Kumar, R. et al. A redox cycle with complex II prioritizes sulfide quinone oxidoreductase-dependent H_2_S oxidation. J Biol Chem 298, 101435 (2022).

27. Blackstone, E. & Roth, M.B. Suspended animation-like state protects mice from lethal hypoxia. Shock 27, 370–372 (2007).

28. Cooper, C.E. & Brown, G.C. The inhibition of mitochondrial cytochrome oxidase by the gases carbon monoxide, nitric oxide, hydrogen cyanide and hydrogen sulfide: chemical mechanism and physiological significance. J Bioenerg Biomembr 40, 533–539 (2008).

29. Petersen, L.C. The effect of inhibitors on the oxygen kinetics of cytochrome c oxidase. Biochim Biophys Acta 460, 299–307 (1977).

30. Titov, D.V. et al. Complementation of mitochondrial electron transport chain by manipulation of the NAD^+^/NADH ratio. Science 352, 231–235 (2016).

31. Volpato, G.P. et al. Inhaled hydrogen sulfide: a rapidly reversible inhibitor of cardiac and metabolic function in the mouse. Anesthesiology 108, 659–668 (2008).

32. Collman, J.P., Ghosh, S., Dey, A. & Decreau, R.A. Using a functional enzyme model to understand the chemistry behind hydrogen sulfide induced hibernation. Proc Natl Acad Sci U S A 106, 22090–22095 (2009).

33. Manford, A.G. et al. A Cellular Mechanism to Detect and Alleviate Reductive Stress. Cell 183, 46–61 e21 (2020).

34. Banerjee, R. & Kumar, R. Gas regulation of complex II reversal via electron shunting to fumarate in the mammalian ETC. Trends Biochem Sci 47, 689–698 (2022).

35. Nicholls, P., Marshall, D.C., Cooper, C.E. & Wilson, M.T. Sulfide inhibition of and metabolism by cytochrome c oxidase. Biochem Soc Trans 41, 1312–1316 (2013).

36. Bostelaar, T. et al. Hydrogen Sulfide Oxidation by Myoglobin. J Am Chem Soc 138, 8476–8488 (2016).

37. Vitvitsky, V. et al. Structural and Mechanistic Insights into Hemoglobin-Catalyzed Hydrogen Sulfide Oxidation and the Fate of Polysulfide Products. J Biol Chem 292, 5584–5592 (2017).

38. Vitvitsky, V., Yadav, P.K., Kurthen, A. & Banerjee, R. Sulfide oxidation by a noncanonical pathway in red blood cells generates thiosulfate and polysulfides. J Biol Chem 290, 8310–8320 (2015).

39. Ruetz, M. et al. A Distal Ligand Mutes the Interaction of Hydrogen Sulfide with Human Neuroglobin. J Biol Chem 292, 6512–6528 (2017).

40. Hill, B.C. et al. Interactions of sulphide and other ligands with cytochrome c oxidase. An electron-paramagnetic-resonance study. Biochem J 224, 591–600 (1984).

41. Lagoutte, E. et al. Oxidation of hydrogen sulfide remains a priority in mammalian cells and causes reverse electron transfer in colonocytes. Biochim Biophys Acta 1797, 1500–1511 (2010).

42. Friederich, M.W. et al. Pathogenic variants in SQOR encoding sulfide:quinone oxidoreductase are a potentially treatable cause of Leigh disease. J Inherit Metab Dis 43, 1024–1036 (2020).

43. Tiranti, V. et al. Loss of ETHE1, a mitochondrial dioxygenase, causes fatal sulfide toxicity in ethylmalonic encephalopathy. Nat Med 15, 200–205 (2009).

44. Di Meo, I. et al. Chronic exposure to sulfide causes accelerated degradation of cytochrome c oxidase in ethylmalonic encephalopathy. Antioxid Redox Signal 15, 353–362 (2011).

45. Brown, G.C., Foxwell, N. & Moncada, S. Transcellular regulation of cell respiration by nitric oxide generated by activated macrophages. FEBS Lett 439, 321–324 (1998).

46. Gnaiger, E. Bioenergetics at low oxygen: dependence of respiration and phosphorylation on oxygen and adenosine diphosphate supply. Respir Physiol 128, 277–297 (2001).

47. Gnaiger, E., Steinlechner-Maran, R., Mendez, G., Eberl, T. & Margreiter, R. Control of mitochondrial and cellular respiration by oxygen. J Bioenerg Biomembr 27, 583–596 (1995).

48. Schindelin, J., et al. Fiji: an open-source platform for biological-image analysis. Nat Methods 9, 676–682 (2012).

