## Supplemental information for "H_2_S preconditioning induces long-lived perturbations in O_2_ metabolism"

Ann Arbor, MI 48109-0600

Corresponding Author: *Address correspondence to: Ruma Banerjee, 4220C MSRB III, 1150 W. Medical Center Dr., University of Michigan, Ann Arbor, MI 48109-0600, email address:

**Figure S1. Sulfide dose-dependent changes in oxygen consumption kinetics. A-B**. HT29 cells exhibit changes in OCR in response to Na_2_S (5-50 µM, red arrow). The black dotted line denotes the basal OCR while the red line indicates the time between sulfide exposure and establishment of a new stationary OCR. **C,D.** Quantitative analysis of data in A, B showing the correlation between Na_2_S dose and recovery time (C), or fractional inhibition of OCR (D). Data are either representative or show mean ± SD of least 3 independent measurements. Two-sample paired *t* test was performed for statistical analysis.

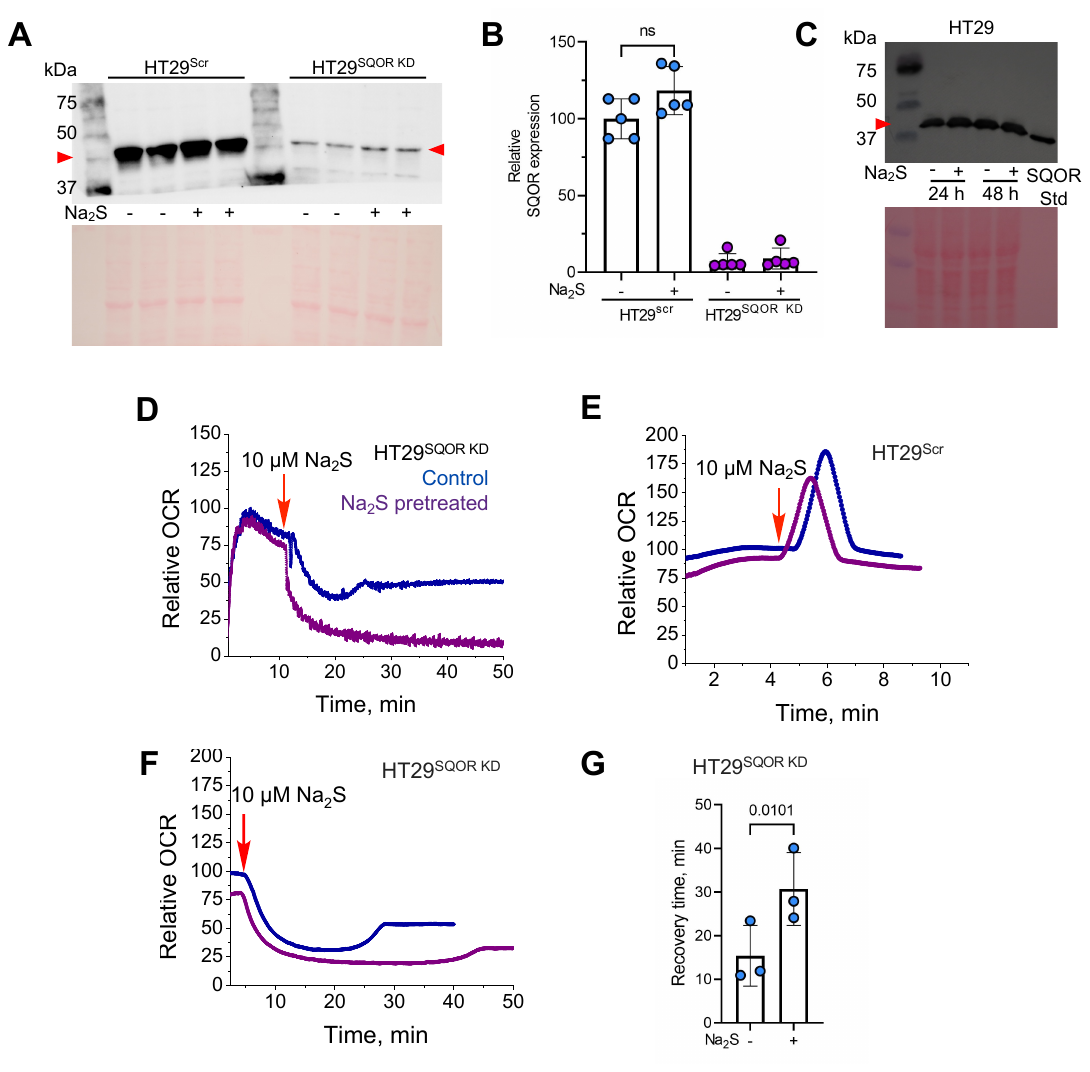

**Figure S2. H_2_S memory is enhanced by SQOR knockdown. A-C**. Western blot analysis showing that SQOR protein levels (red arrowheads) are not affected by sulfide pretreatment (100 µM Na_2_S) in HT29^Scr^ or HT29^SQOR KD^ cells after 4 h (A-B), or in HT29 cells after 24-48 h (C). The lower panels show Ponceau S staining of the membranes as equal loading controls. **D**. Compared to untreated controls (blue), Na_2_S (50 µM, 4 h) pretreatment induced prolonged inhibition of OCR (purple) in HT29^SQOR KD^ cells upon subsequent exposure to 10 µM Na_2_S (red arrow). **E,F**. Compared to untreated controls (blue) Na_2_S (50 µM, 24 h) pretreatment (purple) did not significantly affect OCR in HT29^Scr^ (E) unlike HT29^SQOR KD^ (F) cells, following 10 µM Na_2_S treatment (red arrow). **G**. Na_2_S (50 µM, 24 h) increased recovery time from inhibition by 10 µM Na_2_S in HT29^SQOR KD^ cells. The data are representative of at least 3 independent experiments or show the mean ± SD value (n=3). Two-sample paired *t* test was performed for statistical analysis.

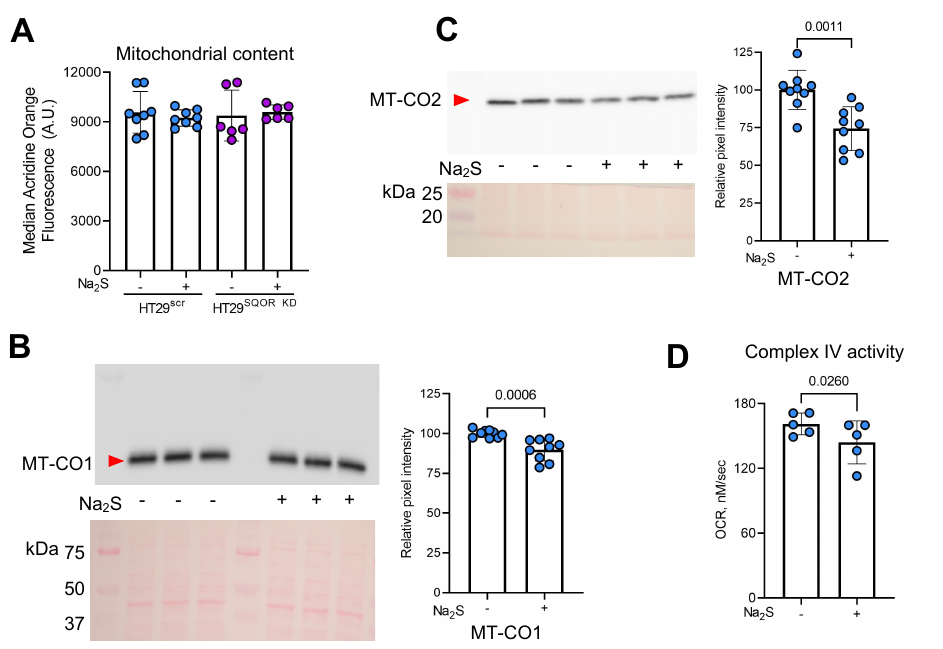
**Figure S3. Sulfide pre-treatment leads to changes in complex IV but not total mitochondria. A.** Na_2_S treatment (100 µM, 24 h) did not change total mitochondrial content as indicated by cardiolipin levels detected with acridine orange staining. (**B,C**) A small but statistically significant decrease in MT-CO1 (E) and MT-CO2 (F) protein levels (upper panels) was observed in response to sulfide pretreatment (100 µM, 4 h). The loading control (lower panels) represents total protein detected by Ponceau S staining. **D**. Complex IV activity as measured in the TMPD assay at 4 h ± 100 µM Na_2_S treatment, was slightly but statistically significantly lower in pretreated cells. The data are representative of 3 independent experiments (B, C) or represent the mean ± SD of at least 5 independent experiments in the remaining panels. Two-sample unpaired (A-C) and paired (D) *t* test was performed for statistical analysis.

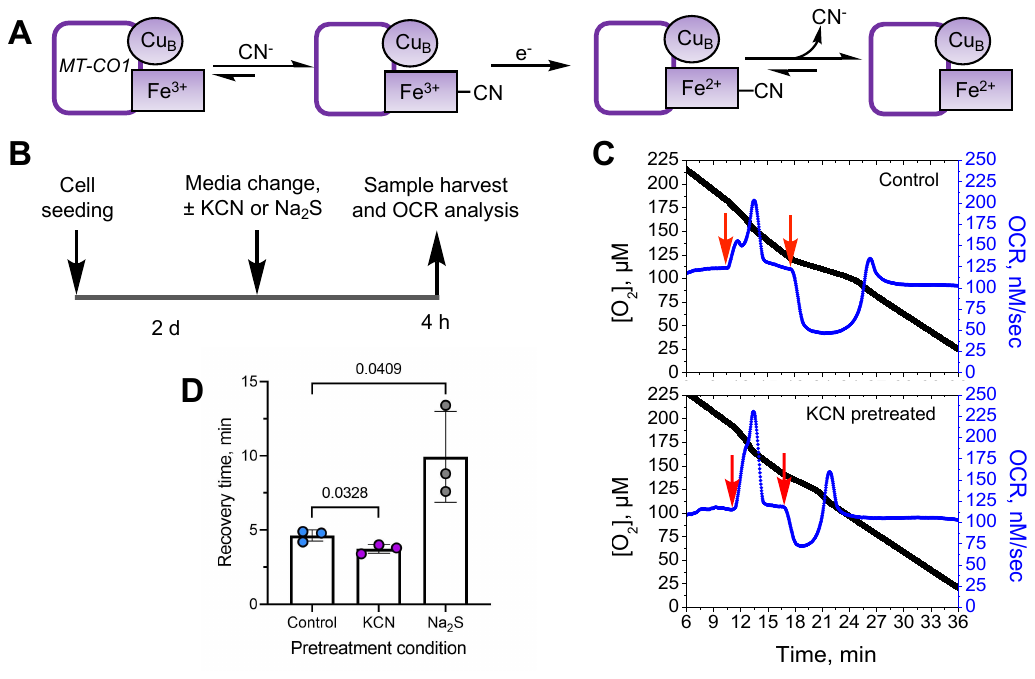

**Figure S4. Acute cyanide exposure does not alter sensitivity to sulfide. A**. Scheme depicting cyanide (CN^-^)-dependent inhibition of complex IV by binding to ferric heme a_3_ in MT-CO1. **B**. Experimental setup used to compare the effects of sulfide (100 µM, 4 h) or cyanide (500 µM, 4h) pretreatment on mitochondrial function. **C,D**. Potassium cyanide (KCN) pretreatment enhanced sulfide-triggered OCR (red arrows) (C) and the recovery time (D) following 20 µM Na_2_S treatment. The data are representative of at least 3 independent experiments (C), and the mean ± SD from these experiments is shown in D. Two-sample unpaired *t* test was performed for statistical analysis.

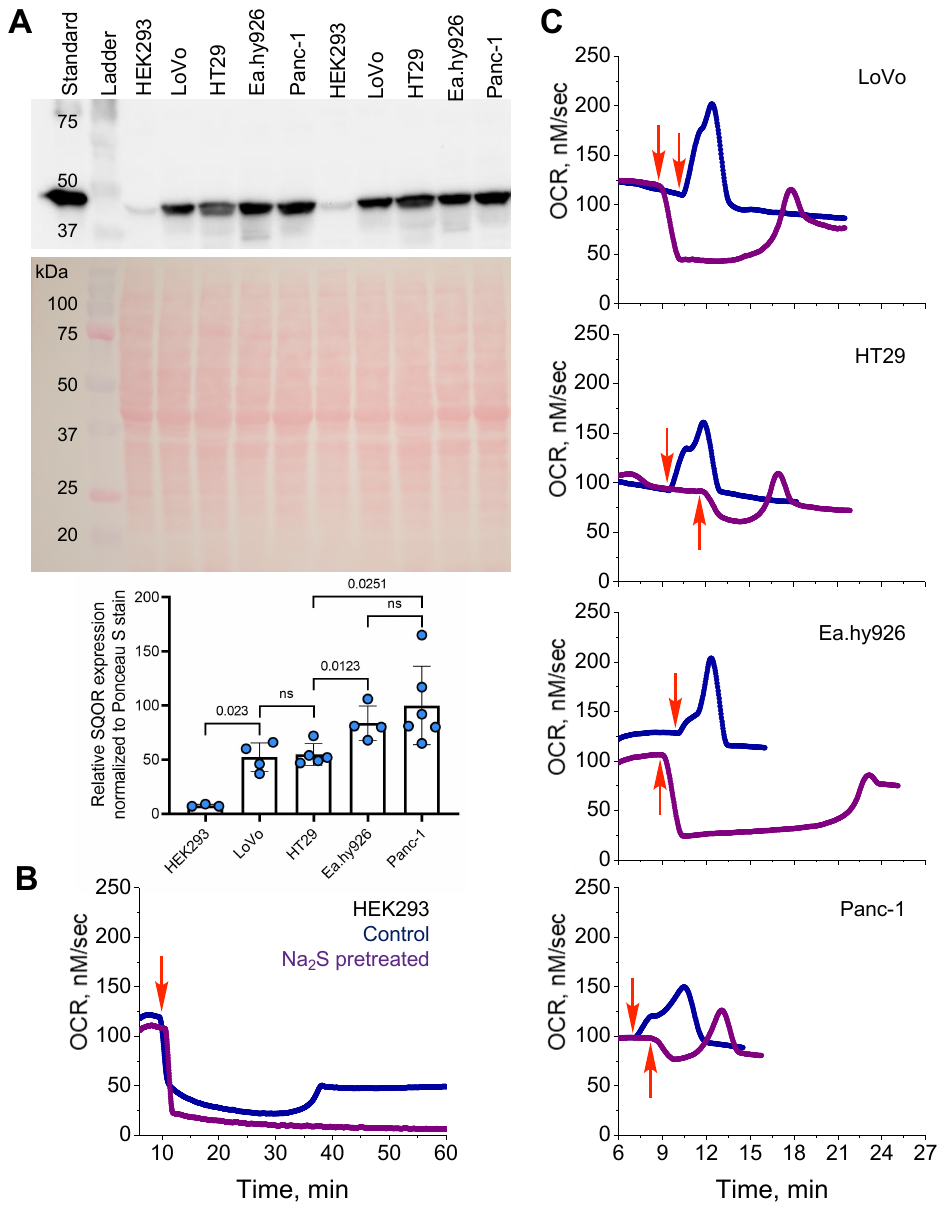

**Figure S5. H_2_S memory correlates with relative SQOR expression in cell lines. A.** SQOR was detected in cell lysates (50 µg protein/lane) from HEK293, LoVo, HT29, Ea.hy926, and Panc-1 cells by western blot analysis (top). Total protein, detected by Ponceau S stain for equal loading (middle) and semi-quantitative analysis of the blot (bottom) are shown. The data are representative of at least 3 independent experiments per cell line. **B,C**. OCR traces for 4 h control (blue) versus 100 µM Na_2_S pretreated (purple) cells following exposure to 20 µM sulfide (red arrow). The data in B-C are representative of 3 independent experiments.

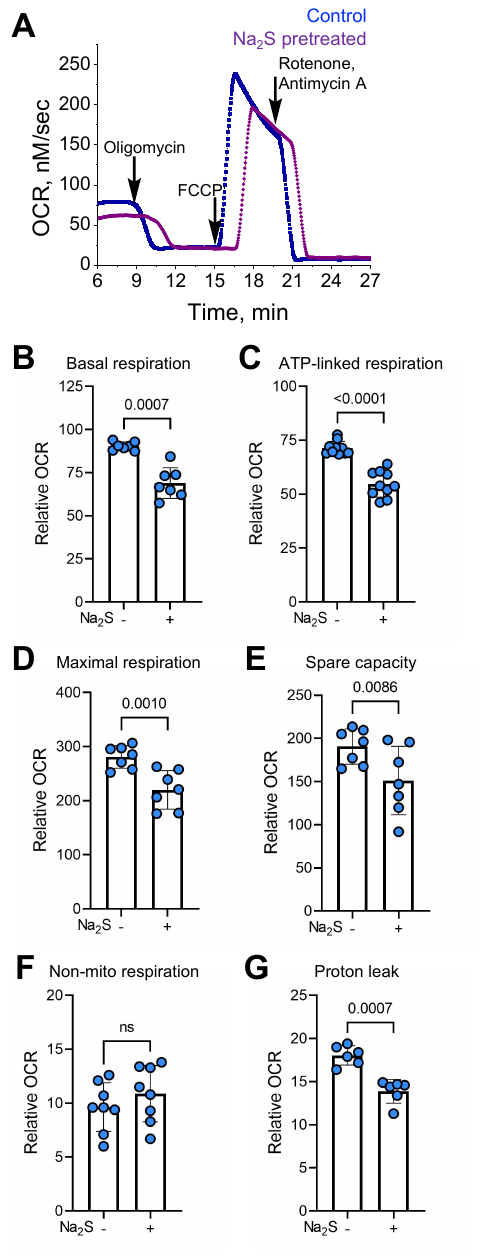

**Figure S6. Acute sulfide treatment alters mitochondrial function. A**. Representative trace of the mitochondrial function profile of HT29 cells 4 h after ± 100 µM Na_2_S treatment in the presence of the following ETC inhibitors: oligomycin (125 nM), FCCP (125 nM), and rotenone and antimycin A (0.5 µM each). **B-G**. Quantitative analysis of the data in A shows the impact of sulfide pretreatment on basal respiration (B), ATP-linked respiration (C), maximal respiration (D), spare capacity (E), non-mitochondrial respiration (F), and the proton leak rate (G). Two-sample paired *t* test was performed for statistical analysis.

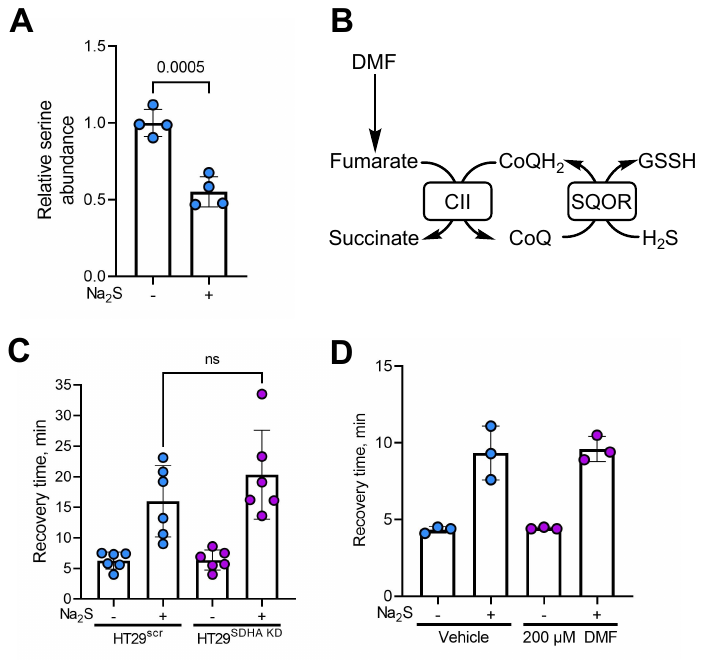

**Figure S7. Sulfide memory is unresponsive to complex II modulation but associated with decreased serine. A.** Sulfide (100 µM, 24 h) exposure leads to a drop in serine levels in HT29 cells. **B.** Scheme showing how complex II (CII) reversal and exogenous dimethylfumarate (DMF) can potentially modulate CoQ availability for SQOR. **C,D.** Knockdown of the SDHA subunit of complex II (C) and 200 µM DMF (D) did not affect recovery time in HT29 cells pretreated with 100 µM Na_2_S for 4 h. Data in C-D are representative of at least 3 independent experiments and are the mean ± SD. Two-sample unpaired (A) and paired (C) *t* test was performed for statistical analysis.
